# Controlled delivery of ultrasound through the head for effective and safe therapies of the brain

**DOI:** 10.1101/2022.12.16.520788

**Authors:** Tom Riis, Matthew Wilson, Jan Kubanek

## Abstract

Transcranial focused ultrasound provides noninvasive and reversible approaches for precise and personalized manipulations of brain circuits, with the potential to transform our understanding of brain function and treatments of brain dysfunction. However, the effectiveness and safety of these approaches have been limited by the human head, which attenuates and distorts ultrasound strongly and unpredictably. To address this lingering barrier, we have developed a “Relative Through-Transmit” (RTT) approach that directly measures and compensates for the attenuation and distortion of a given skull and scalp. We have implemented RTT in hardware and demonstrated that it accurately restores the operator’s intended intensities inside ex-vivo human skulls. Moreover, this functionality enabled effective and intensity-dependent transcranial modulation of nerves and effective release of defined doses of propofol inside the skull. RTT was essential for these new applications of transcranial ultrasound; when not applied, there were no significant differences from sham conditions. Moreover, RTT was safely applied in humans and accounted for all intervening obstacles including hair and ultrasound coupling. This method and hardware unlock the potential of ultrasound-based approaches to provide effective, safe, and reproducible precision therapies of the brain.

---

Transcranial focused low-intensity ultrasound provides a new set of approaches to noninvasively and reversibly manipulate neural activity and treat disorders of brain function.^1,2^. The approaches have included transient^3–5^ and durable^6–11^ modulation of neural circuits, and the delivery of specific drugs across the intact^12–15^ and transiently opened^16,17^ blood-brain barrier. Unlike other noninvasive approaches, ultrasound-based approaches reach millimeter-level precision deep in the brain^18^. Since these approaches are noninvasive and reversible, they provide flexible diagnostic and therapeutic tools that can be applied systematically to address the needs of each individual patient.

However, the effectiveness and safety of these emerging approaches have been hampered by a formidable barrier: the acoustically complex human head. The human skull alone attenuates the ultrasound by a factor of 4.5–64 depending on individual and skull segment ^19–21^. Hair^22,23^, acoustic coupling to the head^24,25^, and entrapped bubbles or air pockets^26^ present additional significant barriers. The joint outcome of these barriers is a severe (**Fig. 1a**) and highly variable (**Fig. 1b**) attenuation^27,28^, which has precluded the delivery of a known ultrasound intensity into desired brain targets. This issue is particularly limiting for emerging reversible therapies, which require the deterministic delivery of effective and safe ultrasound intensity.

**Figure 1.**
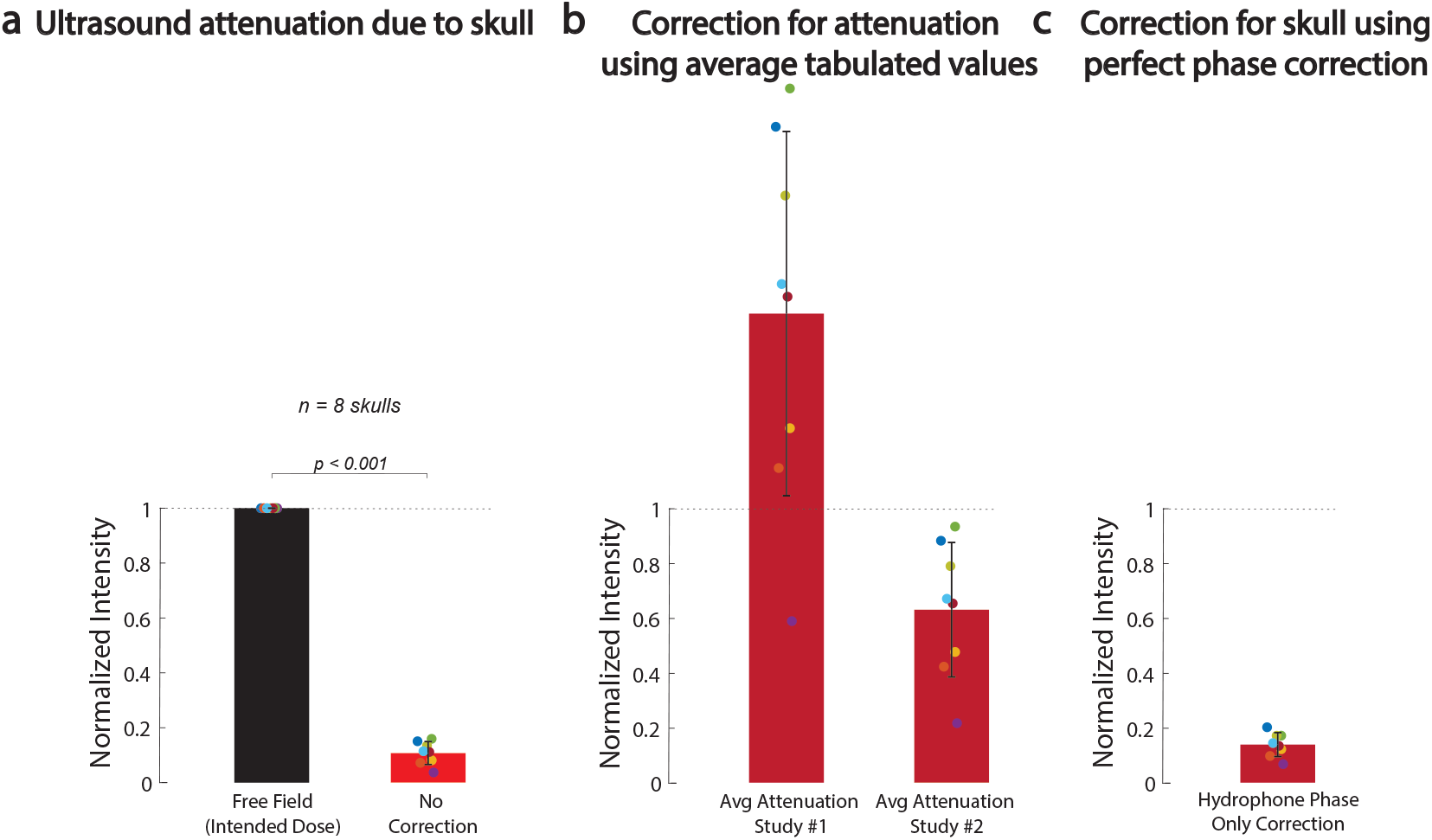
The ultrasound skull attenuation problem. **a,** Spatial peak intensity of the measured field delivered through 8 ex-vivo human skulls into a central target, separately for the intended intensity (black bar) and the intensity following the propagation of ultrasound through the skull (red bar). The intensity field was measured using a calibrated hydrophone (see Methods). **b,** The acoustic attenuation by the skull could be estimated using tabulated values (e.g., study #1^20^ or study #2^21^), but the high variability of the attenuation across individuals makes such estimates inaccurate and uncertain. Same ex-vivo skulls as in **a**. **c,** The problem cannot be addressed using existing methods that correct for the ultrasound dephasing. Same skulls as in panels **a** and **b**, following the hypothetical ideal correction for the ultrasound dephasing using ground-truth hydrophone phase measurements obtained at the target.

The head and the skull additionally dephase each ultrasonic beam. Correction for the phase has been of interest to ablative approaches that aim to maximize the delivered intensity into the brain target^18^. Many methods have been developed to address this specific problem based on CT scans of the head^18,29–31^, ultrasound measurements^32–34^, or other, more invasive approaches^35–39^. Compared with ablations, reversible therapies require the delivery of effective and safe ultrasound intensity. For these reversible applications, the issue of phase aberration is relatively minor compared with the highly variable attenuation by the skull (**Fig. 1c**).

The uncertainties about the intensities delivered into the brain have severely limited emerging reversible applications. This is because these approaches—including neuromodulation and drug delivery—are sensitive to the ultrasound intensity and operate within a narrow window of effectiveness and safety^13,40–42^. Moreover, the lack of knowledge about the intensity delivered into the brain impedes regulatory approval of these emerging reversible therapies.

To address this lingering issue, we have developed an ultrasound-based, “Relative Through-Transmit” (RTT) approach that provides accurate, personalized compensation for the head of each individual and uses the same hardware as that used for the therapeutic intervention. RTT directly measures the attenuation (and dephasing) of a given subject’s head using ultrasound through-transmit measurements, and compensates for it prior to performing the intervention. This method is personalized to each individual and does not require CT, MRI, or other prior information about the head. We implemented RTT in hardware and found that it accurately restores transcranial intensities that operators intend to deliver into the brain. We show that this capacity is critical for effective modulation of excitable structures and effective drug release in deep brain locations. Moreover, RTT can be safely and practically applied through the human head.

## Results

**Fig. 1a** demonstrates the severity of acoustic attenuation by the skull. Across 8 *ex-vivo* skulls, we found that the ultrasound intensity delivered into a deep brain location (see Methods) is attenuated by a factor of 11.4± 6.8 (mean±S.D.), which replicates previous findings^43^. In principle, the attenuation could be estimated using tabulated values (e.g.,^20,21^), but the high variability of the attenuation across individuals makes such estimates inaccurate and uncertain (**Fig. 1b**). For instance, using the values of those two studies would over- and under-estimate the average value by a factor of 1.7 ± 0.67 and 0.63 ± 0.25, respectively, and lead to high variability (pooled standard deviation equal to 0.47, compared to the normalized intensity of 1.0). A compensation for the dephasing of the ultrasound, which can be obtained using existing methods^18,29,30,32–39^ is useful for ultrasound-based surgeries^18^ and to an extent also for the present purpose of delivering deterministic intensity into specified targets for repeated applications (**Fig. 1c**). Nonetheless, even the hypothetically ideal correction for the phase based on ground-truth measurements (**Fig. 1c**) leaves an average discrepancy of 85% between the intended and actual intensities delivered into a brain target.

### RTT and hardware

RTT rests on two sets of transducer phased arrays positioned at opposite sides of the head (**Fig. 2**). Each element of the array operates in both transmit and receive modes. We implemented this geometry using two sets of 128-element arrays, and connected the arrays to a driving system that can independently transmit signals from and listen to each element. This complete through-transmit system enables ultrasound-based correction for the attenuation (and dephasing) of each ultrasonic beam (**Fig. 2**).

**Figure 2.**
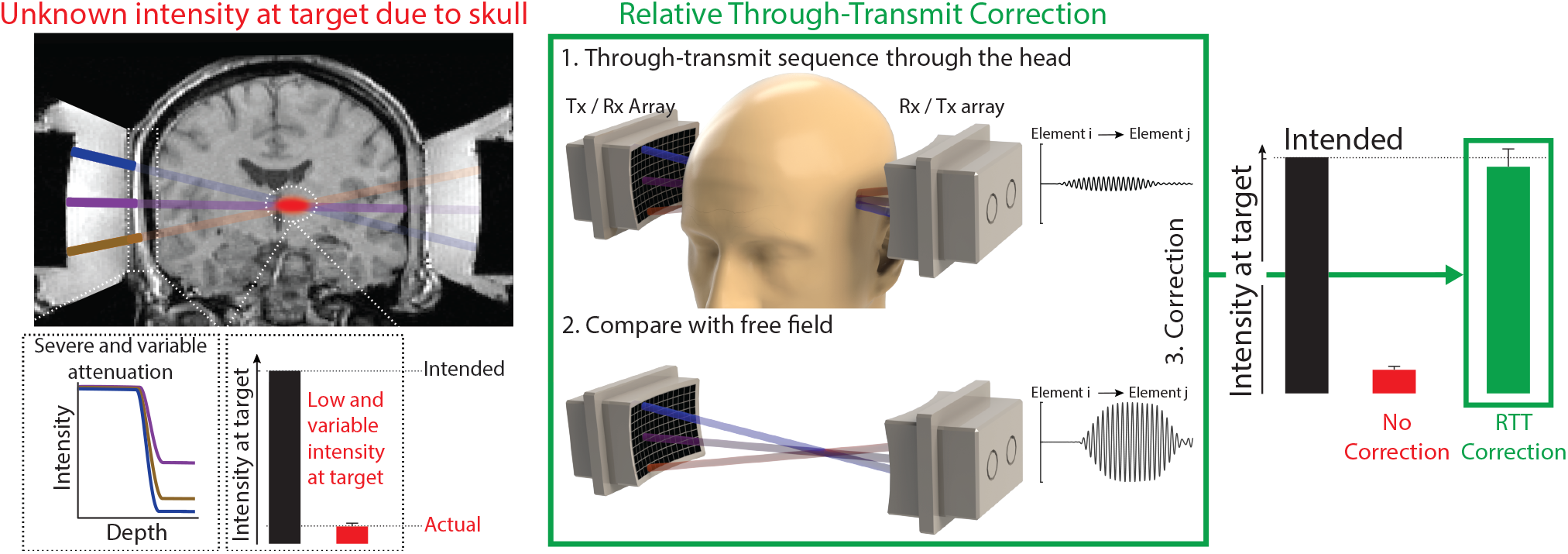
Method for controlled delivery of ultrasound into the brain. Problem (red): The human skull precludes the delivery of controlled ultrasound dose into the brain. The human skull attenuates ultrasound strongly and unpredictably, leading to low and variable intensity delivered into a brain target. This is illustrated by the red target inside an MRI scan of a human subject and the red bar representing the actual intensity electronically focused into a central target through an *ex-vivo* skull. **Solution (green).** We developed a method, RTT, which uses brief through-transmit pulses of low-intensity ultrasound to measure the ultrasound aberrations by the skull. The top waveform is an example of a typical through-transmit signal recorded through the head of a participant with unshaved hair. The scan is performed with a subject’s head in the ultrasound path, and the obtained signals are compared with the signals received through water (bottom signal). From the relative differences in the magnitudes and times of flight of the received signals, RTT computes the attenuation and phase shift of each segment of the skull within each ultrasonic beam. These values are then used to scale up and delay the firing of ultrasound from the individual elements and thus effectively compensate for the skull, restoring the intended intensity at the target (right bars; green). No CT or MRI images of the head are required. The green bar shows the mean ± s.e.m. correction value across 8 ex-vivo skulls and 3 targets (see further below). The details of the scan and the computation are provided in the Methods.

### Transcranial delivery of deterministic intensities

To test the accuracy of RTT, we used the same hardware to electronically focus ultrasound into specific targets inside human *ex-vivo* skulls. We measured the induced fields using a hydrophone (see Methods). We evaluated the measured intensities in four conditions. First, we measured the intensities in free-field, which presents no obstacles for ultrasound. This intensity corresponds to the intensity intended to be delivered into the target by the operator. Second, we introduced skulls between the hardware and the hydrophone, and measured the resulting intensities. This case presents the worst-case scenario of no correction for the skull. Third, we evaluated the best possible scenario, the hypothetical ideal correction for the skull. To do that, we used the hydrophone to measure the attenuation and dephasing for each element of the device, thus obtaining ground-truth values. We used these ground-truth values to compensate for these aberrations, scaling the magnitude of the emitted ultrasound from each element and delaying it accordingly (see Methods for details), as if no skull was present. And fourth, we applied the RTT correction.

We performed these measurement inside 8 water-immersed, degassed human *ex-vivo* skulls. **Fig. 3** shows the spatial peak intensity and the associated field for a target positioned at the center of the two transducers. The figure reinforces the notion that human skulls severely dampen the intensity delivered into the brain (red). Compared with the free-field values, the ultrasound intensity through the skull was attenuated by a factor of 11.4 ± 6.8 (mean±S.D.), degrading it to 10.7 ± 4.2% of the intended intensity. The difference between the free-field and through-skull values was significant (*t*_7_ = 60.7, *p* = 8.6 × 10^−11^, paired two-tailed t-test).

**Figure 3.**
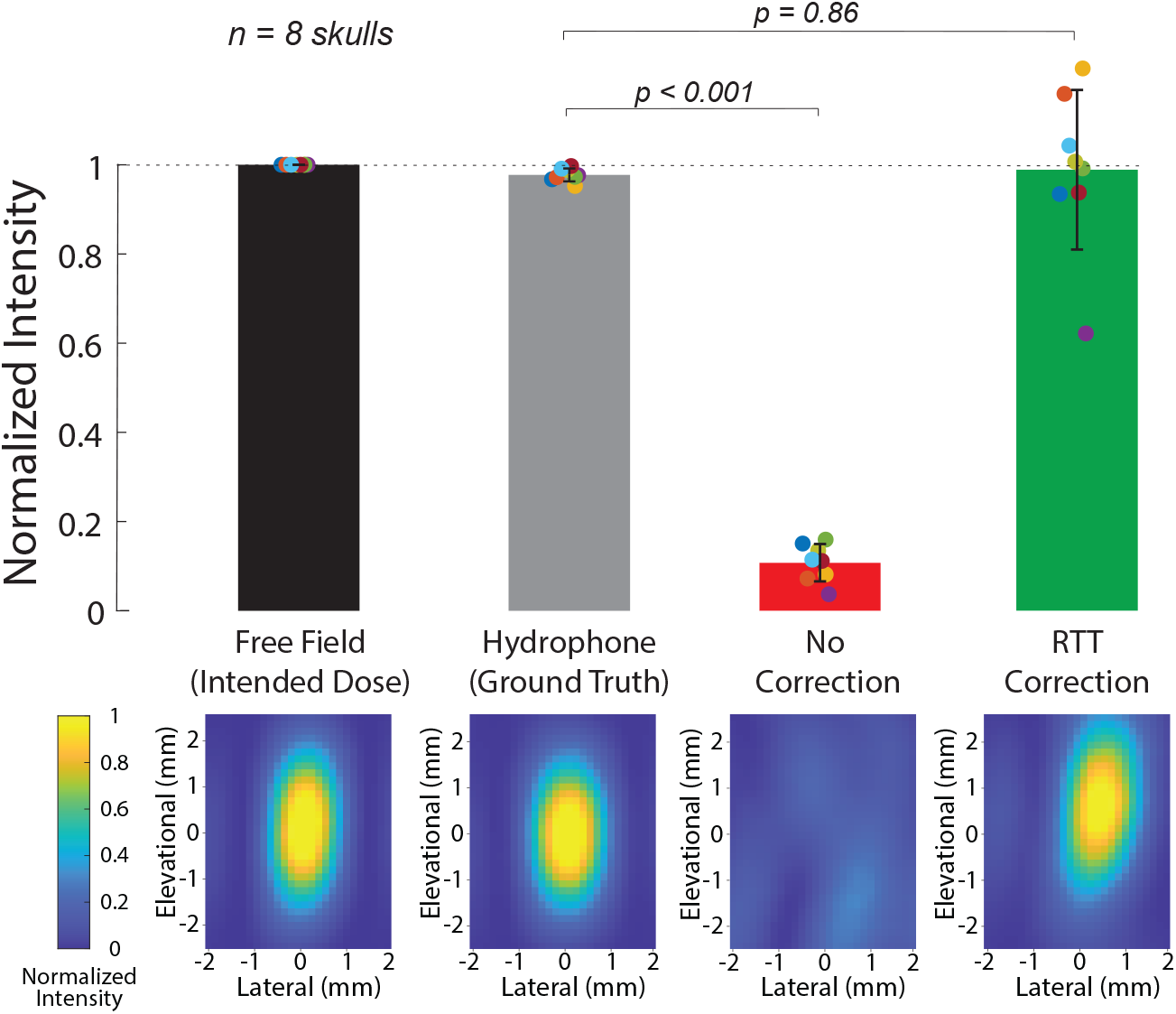
RTT accurately compensates for each skull and restores the intended intensity at target. Ultrasound fields obtained inside ex-vivo human skulls (*n* = 8), separately for the hypothetical ideal correction (gray), no correction (red), and RTT (green). The top bars show the spatial peak intensities of the field for each case. The bottom plots provide the corresponding spatial distribution of the ultrasonic fields with respect to the target.

We then applied RTT using the phased arrays. **Fig. 3** shows that RTT restores the intended intensity values (green). The RTT-compensated intensity constituted 98.8±17.8% (mean±S.D.) of the intended values in free-field, and there was no significant difference between the mean of two conditions (*t*_7_ = 0.18, *p* = 0.86, paired two-tailed t-test). The average value was also not significantly different than the hypothetical, best-possible correction based on the hydrophone ground-truth measurements inside the skull (black bar; *t*_7_ = 0.17, *p* = 0.86, paired two-tailed t-test). One skull (purple datapoint) attenuated the ultrasound severely (a factor of 26.9 attenuation). This was likely be due to visually present outgrowths possibly related to hyperostosis, as assessed by a neurosurgeon. For this skull, the RTT correction was less accurate, achieving a factor of 0.62 of the intended intensity.

We next assessed whether RTT could be applied to the human head, which presents additional key barriers for transcranial ultrasound including hair, scalp, acoustic coupling, and the brain. This test also evaluated the safety of the method. RTT was designed to be safe. The RTT scan consists of brief (< 100*μ*s) low-intensity (average peak pressure of 80 kPa in free field; **Suppl. Fig. 4**) pulses of ultrasound. The RTT scan takes less than one second to complete. Subjects (*n* = 5) did not feel any discomfort during the procedure. **Fig. 4**-blue shows the average through transmit attenuation through both sides of the head, separately for each subject. The figure demonstrates that the method offers through-transmit quality comparable to the ex-vivo skulls (gray). Specifically, the receiving elements on the opposite side of the head recorded (**Fig. 2**) an average of 7.3 ± 4.8% (mean ± SD, *n* = 5 subjects) of the signal amplitude when RTT was applied through the human skull, and 12.6 ± 8.2% for the characterized ex-vivo human skulls (mean ± SD, *n* = 8 skulls). The additional factor of 1.7 attenuation is expected because the application of ultrasound through the human head incurs additional attenuation by hair, scalp, coupling, bubbles or air pockets in between, as well as tissues inside the skull. No subject reported side effects at 1 week follow-up. Thus, RTT can be safely applied to the head of humans and directly measures the attenuation by all obstacles within the ultrasound path.

**Figure 4.**
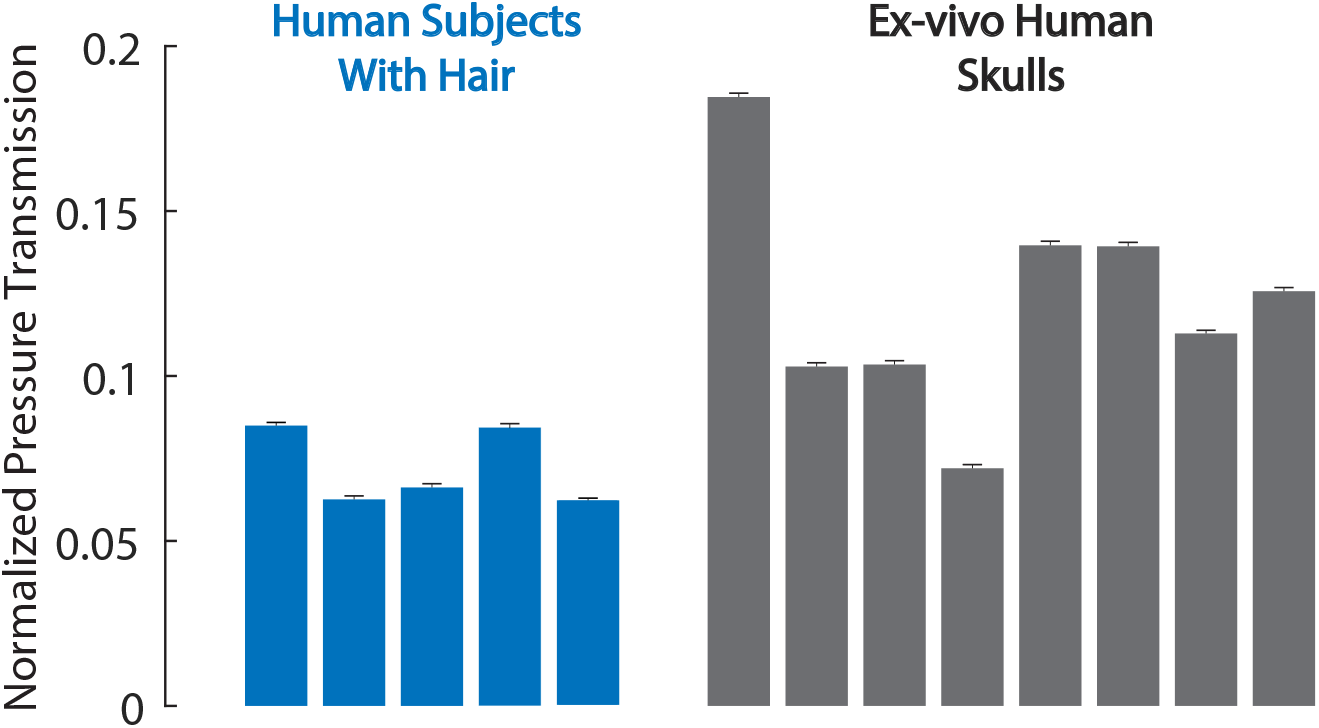
RTT applied though the human head. Average through transmit attenuation value across all elements in 5 human subjects and 8 *ex-vivo* human skulls. No hair shaving was necessary to obtain robust through-transmit signals (see also **Suppl. Fig. 8**).

We next tested the robustness of RTT with respect to brain target location. To do so, we used the phased arrays to refocus the ultrasound into targets covering the full steering range of the device: 10 mm axial, 20 mm axial, 10 mm lateral, 20 mm lateral, and 15 mm elevational to the central target (**Fig. 5**). RTT correction brought the delivered intensity to 96.3 ± 21.4%, 94.8 ± 23.2%, 92.8 ± 16.4%, 62.5 ± 15.7%, and71.6 ± 18.03% of the intended value in each target respectively. There was no statistical difference between the mean of intended peak intensities and the RTT-compensated peak intensities at the central target (*t*_7_ = 0.18, *p* = 0.86, paired two-tailed t-test), 10 mm axial (*t*_7_ = 0.48, *p* = 0.64, paired two-tailed t-test), 10 mm lateral (*t*_7_ = 0.42, *p* = 0.55, paired two-tailed t-test), and 20 mm axial (*t*_7_ = 1.23, *p* = 0.26, paired two-tailed t-test). There was a significant difference in the average delivered intensity at the target 20 mm lateral (*t*_7_ = 6.748, *p* = 0.0002, paired two-tailed t-test) and 15 mm elevational (*t*_7_ = 4.5, *p* = 0.003, paired two-tailed t-test).

**Figure 5.**
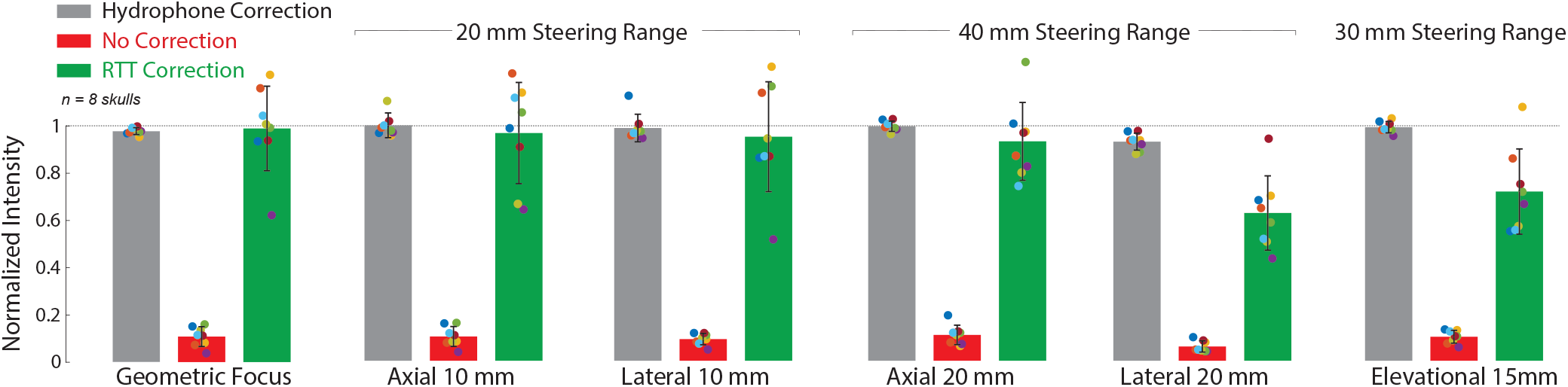
RTT performance as a function of the range of action. Same format as in **Fig. 3**, now also including targets comprising the full steering range of the array: 10 mm axial, 20 mm axial, 10 mm lateral, 20 mm lateral, and 15 mm elevational to the central target. Axial refers to the line connecting the centers of the two transducers.

We subsequently tested the relative contribution of the two key components of the ultrasound aberration by the skull—the attenuation and dephasing. **Fig. 6** shows the spatial peak intensity following the engagement of each correction type in isolation as well as their joint application. At the central target, phase-only correction resulted in average intensity of 13.8 ± 4.3% (mean±S.D.) for the ideal hydrophone correction (gray) and 11.9 ± 4.9% for RTT (green). The no correction value (red) was 10.7 ± 4.2% of the free field intensity. The amplitude-only correction brought the peak spatial intensity to 71 ± 12.5 for the hydrophone and 93.8 ± 28.9 for RTT. Thus, the correction for the attenuation (i.e., for the amplitude of the received signals) constitutes the key factor in the delivered ultrasound intensity. The inclusion of the correction for the phase is additionally desirable in that the resulting joint correction (**Fig. 6c**) brings the average delivered intensity to 98.8±17.8%.

**Figure 6.**
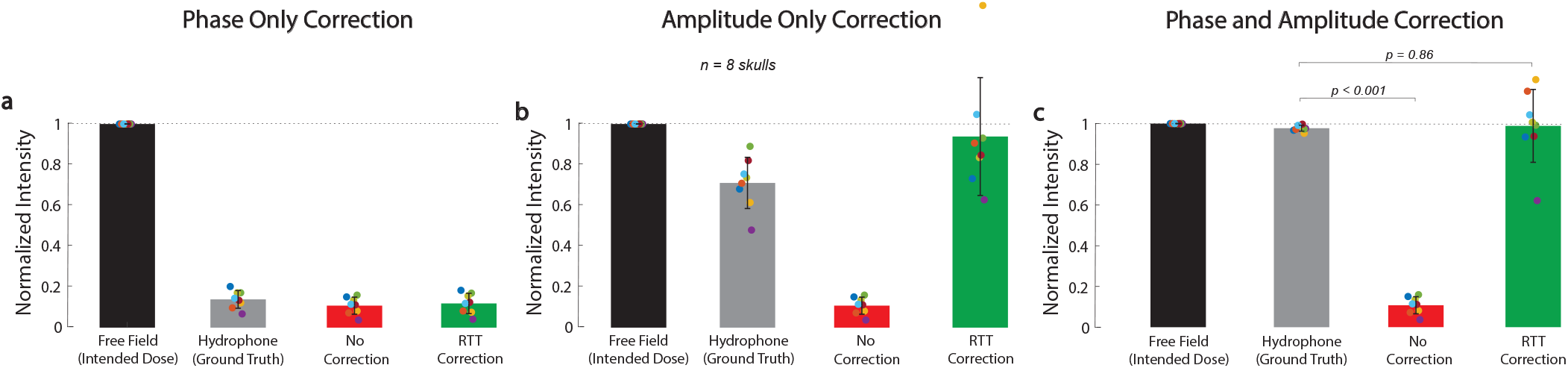
Correction for phase is insufficient to account for the skull attenuation. Spatial peak intensity at the central target using the phase correction component (**a**), amplitude component (**b**) and both components (**c**) of RTT. Same format as in **Fig. 3**.

We further tested the robustness of RTT with respect to specific hardware. In particular, we additionally implemented RTT on arrays that had the same number of elements but much larger aperture (**Suppl. Fig. 1**). For this configuration, skulls (*n* = 4 specimens) degraded the intensity at the geometric center to 6.3 ± 1.7% of the intended, free-field value, in line with **Fig. 1**. The RTT compensation recovered the intensity at the target to 104 ± 18.1% of the intended value. Following the compensation, there was no significant difference between the intended and mean RTT-recovered intensities (*t*_3_ = 0.47, *p* = 0.67, paired two-tailed t-test).

We evaluated the clinical utility of the approach to two emerging applications of transcranial ultrasound—neuromodulation and drug release.

First, we tested how ultrasound could be applied through the skull to stimulate nerves. To test effects on nerves within intact biological tissues, we instructed 11 human subjects to place their thumb into a holder at the central target inside an ev-vivo skull. We quantified the subjects’ responsiveness to the ultrasound when RTT was applied and when it was absent (see Methods). We delivered into the target a 300 ms stimulus of specific pressure levels and assessed the effects on the subjects’ nociceptive responses. Nociceptive responses indicate stimulation of nerves or nerve endings in the tissue^5,44,45^. We found that RTT was critical for effective stimulation (**Fig. 7a**). Without RTT, there was no significant stimulation (red; *t*_11_ = 1.00, *p* = 0.34, one-sample two-tailed t-test). Following RTT, the response rate of subjects to the stimuli reached 62.7%. This level was statistically equivalent (*t*_10_ = 0.58, *p* = 0.57, paired two-tailed t-test) to a 66.3% response rate obtained with the hypothetical best-possible ground-truth correction, which is as good as if no skull was present. **Suppl. Fig. 3** show individual responses to each correction of all subjects.

**Figure 7.**
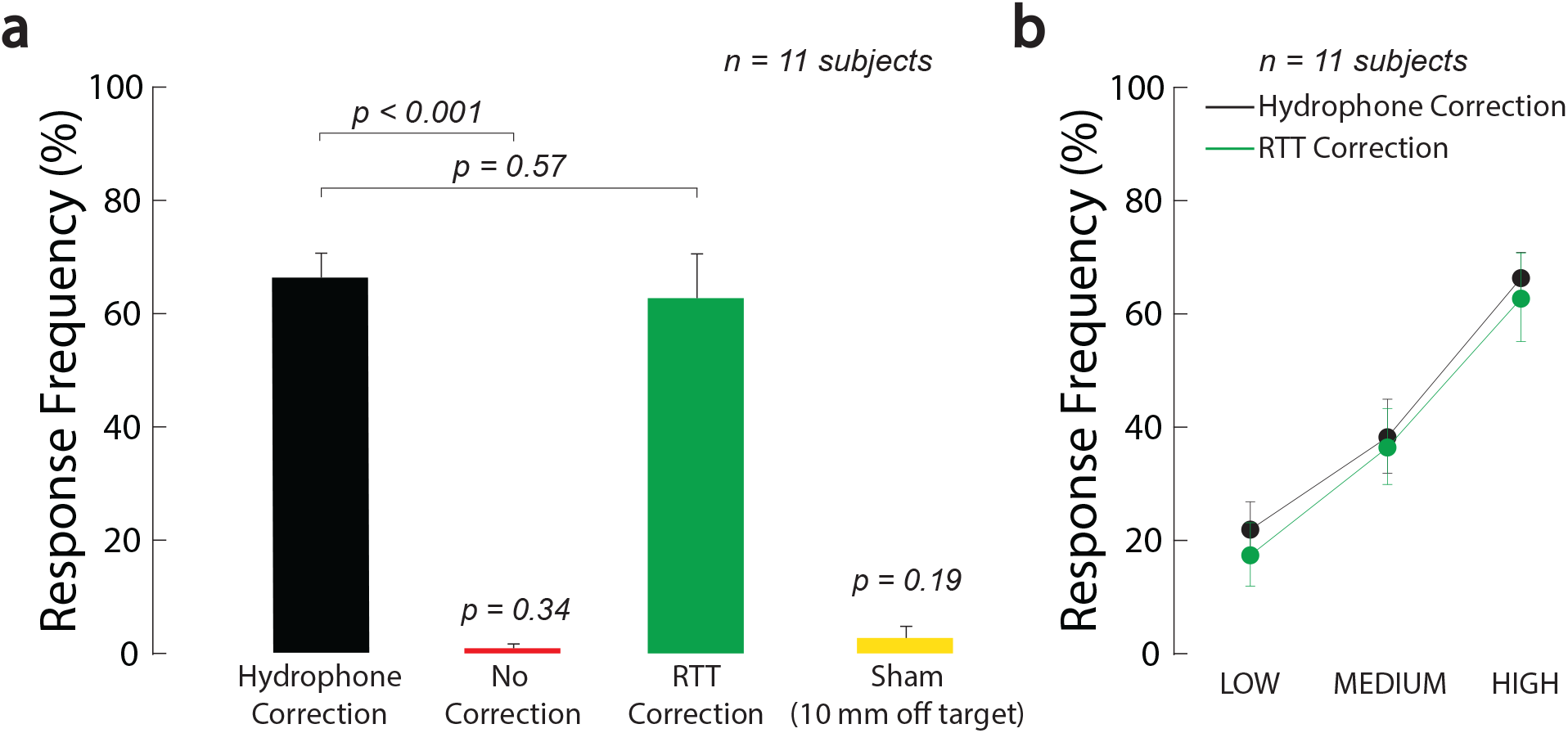
RTT enables effective ultrasonic stimulation through the skull. **a, RTT enables effective modulation of peripheral nerves through skull.** The arrays targeted nerves in the thumb of 11 participants. The thumb was secured in the central target inside an ex-vivo skull. The arrays delivered into the target a 300 ms stimulus at a frequency of 650 kHz and pressure amplitude of 1.8 MPa. The data were collected with the ideal correction (black), without any correction (red), and after applying RTT (green). A sham condition delivered the stimulus 10 mm below the finger (yellow). The individual conditions were presented randomly every 8-12 s, for a total of 10 repetitions. Subjects reported any nociceptive response, which indicates stimulation of nerves and nerve endings. Response frequency represents the proportion of trials in which subjects reported a nociceptive response. **b, Dose-response relationship of the stimulation.** There was a significant modulation by the ultrasound pressure but no significant difference in the responses following the ideal (black) and RTT (green) corrections (see text for details). The LOW, MEDIUM, and HIGH labels correspond to an intended peak pressure of 1.3, 1.55, and 1.8 MPa, as measured in free-field. The error bars represent the s.e.m.

To control for potential confounds that could be associated with ultrasonic stimulation, we randomly interleaved a sham stimulus that delivered the ultrasound 10 mm below the target with hydrophone correction. This off-target stimulation produced no significant stimulation (yellow, *p* = 0.19, one-sample two-tailed t-test, *t*_11_ = 1.39). This controls for a potential artifactual effect and confirms the spatial specificity of the stimulation.

We further investigated the dose dependence of the stimulatory effects. Specifically, we varied the stimulation across three intensity levels, all within the FDA 510(k) Track 3 guidelines^42^. We found an increase in stimulation effectiveness with increasing level of the ultrasound (**Fig. 7b**). The response frequency reached 62.7% for the strongest (1.8 MPa) stimulus, and was significant also for the weakest stimulus tested (1.3 MPa; *t*_10_ = 7.63, *p* = 1.7 × 10^−5^, one-sample two-tailed t-test). The effect of the stimulation level was highly significant (two-way ANOVA, *F*_2,60_ = 25.24, *p* = 1.1 × 10^−8^). The responses were statistically indistinguishable from the hypothetical ideal correction (green versus black; two-way ANOVA, *F*_1,60_ = 0.41, *p* = 0.52), and there was no significant interaction between the two factors (*F*_2,60_ = 0.20, *p* = 0.98).

Therefore, the accurate compensation for the delivered intensities into brain targets (**Fig. 3**, **Fig. 5**) also translates into restored stimulation effectiveness with response levels not attainable without this method (**Fig. 7**).

Second, we tested whether RTT could be used to release drugs at clinically-relevant and deterministic doses in specific locations inside the skull. To do so, we devised ultrasound-sensitive nanoparticle carriers^12,46,47^, and encapsulated in the nanoparticles the neuromodulatory drug propofol at a concentration of 0.063 mg/ml (Methods). We then tested how the nanoparticles respond to ultrasound when RTT correction is applied and when it is not, in a manner analogous to **Fig. 7**. We found that RTT is critical to mediate effective release when ultrasound is applied through the skull (**Fig. 8a**). Without RTT (red), the amount of detected drug was no different (*t*_14_ = 0.30, *p* = 0.77, two-sample two-tailed t-test) from the no stimulation case (purple). The application of RTT (green) nearly tripled the release effectiveness (factor of 2.9 increase), releasing 31.6% of the encapsulated propofol. This level was statistically equivalent (*t*_18_ = 0.08, *p* = 0.94, paired two-tailed t-test) to the 31.8% release obtained with the hypothetical best-possible correction (black).

**Figure 8.**
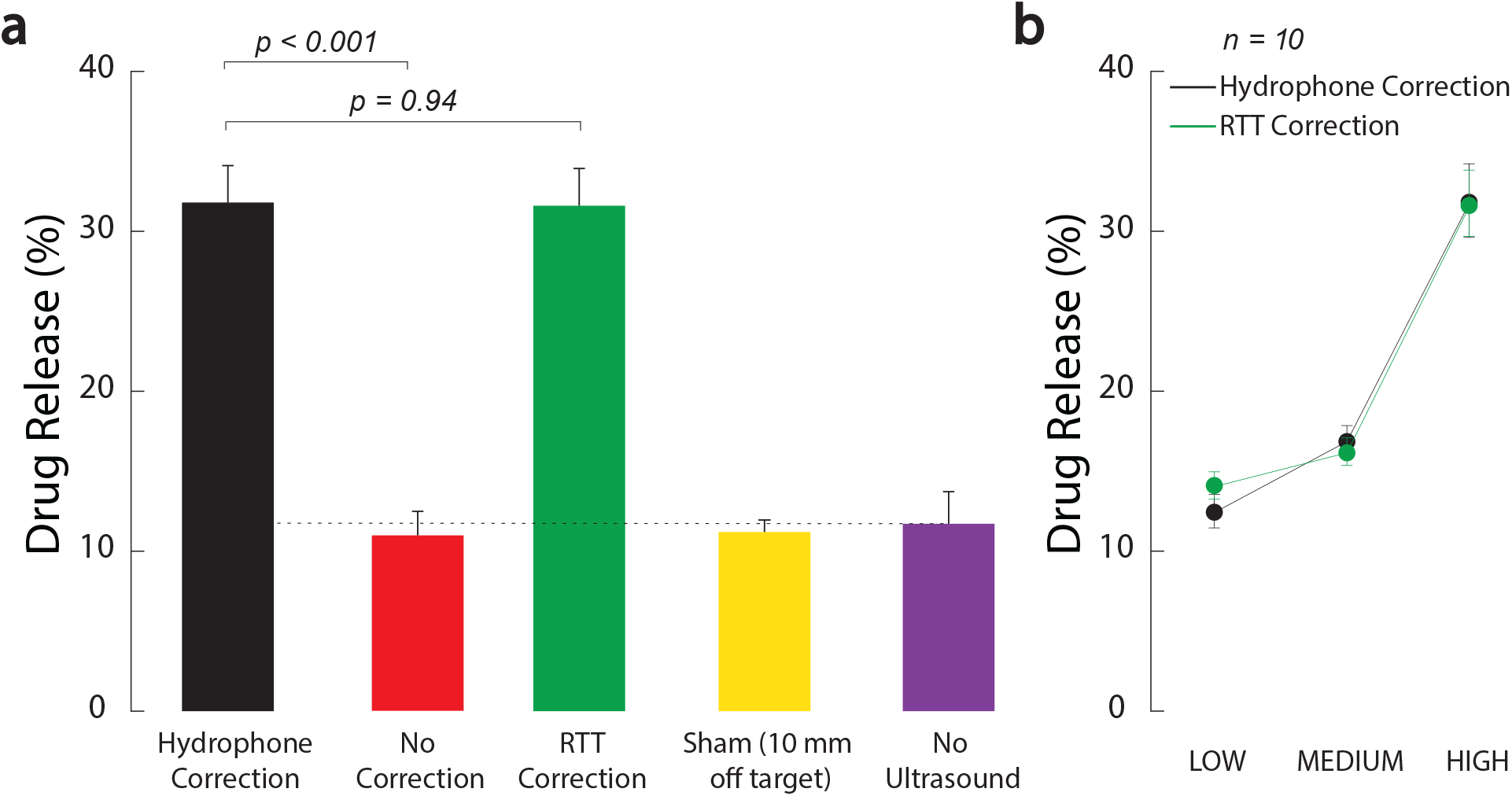
RTT enables effective and dose-dependent local drug release. **a, RTT enables effective drug release from nanoparticle carriers.** Safe, biocompatible nanoparticle carriers^47^ encapsulated the neuromodulatory drug propofol. The nanoparticles release their drug load when impacted by low to medium intensity ultrasound. Vials with the nanoparticles were positioned within a central location of an *ex-vivo* skull, analogously to **Fig. 7**. Ultrasound, delivered through the skull, impacted the nanoparticles in 100 ms pulses delivered every 1 s for 60 s, at a frequency of 650 kHz and pressure amplitude of 1.8 MPa. The data were collected under the hypothetical ideal correction for the skull (black), without any correction (red), and after applying RTT (green). A sham condition delivered the stimulus 10 mm below the vial (yellow). A second sham condition placed the vial at target but delivered no ultrasound (purple). The dotted line represents the baseline when no ultrasound is applied. The baseline can be non-zero due to free (unencapsulated) drug or due to the nanoparticles being partially leaky^47^. Analogously to **Fig. 7**, the individual conditions were randomly interleaved. The bars comprise *n* = 10 distinct samples, with the exception of the No Ultrasound case, which used *n* = 6. The error bars represent the s.e.m. **b, Dose-response relationship.** The LOW, MEDIUM, and HIGH labels correspond to an intended peak pressure of 1.2, 1.5, and 1.8 MPa, as measured in free-field. All datapoints comprised *n* = 10 distinct samples. The error bars represent the s.e.m.

We confirmed the spatial specificity of the release using a sham condition in which the ultrasound was focused 10 mm below each vial (**Fig. 8a**, yellow bar). In this case, the amount of detected drug was no different (*t*_14_ = 0.30, *p* = 0.77) from the case in which no ultrasound was applied (purple).

We finally investigated the dose dependence of the release. To do so, we varied the delivered ultrasound intensity across the same levels as in **Fig. 7**. We found an increase in stimulation effectiveness with increasing level of the ultrasound (**Fig. 8b**). The effect of the stimulation level was highly significant (two-way ANOVA, *F*_2,54_ = 84.53, *p* = 2.3 × 10^−17^). The release levels were statistically indistinguishable from the hypothetical ideal correction (green versus black; two-way ANOVA, *F*_1,54_ = 0.02, *p* = 0.89), and there was no significant interaction between the two factors (*F*_2,54_ = 0.35, *p* = 0.7).

Therefore, RTT can be used to release a controlled dose of drugs in spatially localized regions within the human skull.

## Discussion

The human skull has been a formidable barrier for current and emerging applications of ultrasound to the brain. This barrier has precluded effective, safe, and reproducible neurointerventions based on low-intensity ultrasound. This issue has been limiting in that effectiveness *and safety* are critical premises for next-generation reversible and repair therapies of the brain.

We developed an ultrasound-based method, RTT, that compensates for the strong and variable attenuation of ultrasound by the skull (**Fig. 3**, **Fig. 6**, **Fig. 7**). In addition to attenuation, RTT also compensates for the dephasing of ultrasound by the skull, although the dephasing was found to be a relatively minor issue (**Fig. 6**) and can be satisfactorily performed using existing CT-based approaches^18,29–31^. We implemented RTT in hardware and showed that it faithfully restores the intensities delivered into individual brain targets. The deterministic delivery of ultrasound through the skull enabled effective transcranial stimulation of intact peripheral nerves and the release of deterministic amounts of a neuromodulatory drug from nanoparticle carriers. RTT was found to be robust with respect to treatment location, ultrasound intensity levels, and two distinct hardware embodiments. Finally, RTT was safely applied to the human head (**Fig. 4**).

The delivery of deterministic ultrasound intensity through the skull (**Fig. 3**) opens the path to all applications of transcranial ultrasound which aim to deliver effective yet reversible therapies. We have shown example applications of RTT to modulation of peripheral nerves and to release of propofol from nanoparticle carriers (**Fig. 7**, **Fig. 8**). RTT was key for effective neuromodulation and drug release (**Fig. 7a**, **Fig. 8a**) and also for their deterministic dosing (**Fig. 7b**, **Fig. 8b**).

The deterministic dosing is crucial with respect to effectiveness and safety. For instance, nanoparticle carriers require relatively high ultrasound pressures for effective release—1.8 MPa (**Fig. 8b**), which reproduces findings of previous studies^13,47^. This level lies close to the FDA *I*_SPPA_ 510(k) Track 3 limit (*I*_SPPA_ < 190 W/cm^242^, corresponding to about 2.4 MPa in soft tissues). Therefore, there is a narrow window between effectiveness and safety. This narrow window requires an accurate compensation for the skull. Microbubble-mediated opening of the blood-brain barrier^16,48^ is even more sensitive to the ultrasound exposure. For example, a difference between 0.7 MPa and 0.8 MPa dictates whether blood-brain barrier in the human brain remains intact or is opened^48^. RTT is therefore expected to accelerate the regulatory approval of these emerging applications.

RTT can be safely applied to the human head. We have applied RTT to five human participants with hair (**Fig. 4**). No subject experienced detrimental effects during or following the procedure. This result is expected because RTT uses diagnostic-imaging-like, < 100 *μ*s pulses that were safely within the FDA 510(k) Track 3 guidelines^42^: *I*_SPPA_ = 1.3 W/cm^2^, *I*_SPTA_ = 5.4 mW/cm^2^. The subjects did not perceive heating, which is also as expected given the low RTT energies. Coupling was provided using a biocompatible hydrogel^49^. To effectively target specific brain regions, the MRI compatible transducers can be registered to the brain target using MRI (**Fig. 2**). MRI-free operation may also be possible using standard optical tracking devices^50^.

The compensation for the skull could conceivably heat the skull during the ensuing neuromodulation or other low-intensity applications. For common low-intensity applications, this is, however, unlikely. Specifically, the average pressure transmission per element across all skulls and targets measured with RTT was 38 ± 19%, corresponding to an average pressure amplitude scaling by 3.41 ± 2.15. Even following this scaling, the overall energy deposited into the skull for low intensity BBB and neuromodulation therapies (i.e., 0.5 MPa for 300 ms) is several orders of magnitude lower than the exposures that can produce harmful skull heating^51^. Nonetheless, any new application of transcranial ultrasound to the brain should proceed in a staged manner, with intensity being increased gradually while carefully monitoring safety of the brain and the skull.

For the neuromodulatory application tested here, we used pulses of frequencies in the neuromodulatory and drug release range^52^. For any application, the frequency can be set to match that used for subsequent treatments, thus providing a direct measurement of the skull aberrations for that particular frequency.

RTT compensates for both the ultrasound attenuation by the skull (critical; **Fig. 6**) and the ultrasound dephasing (beneficial; **Fig. 6**). Current FDA-approved treatments use high-intensity ultrasound to perform ablative surgeries^18^. For these applications, it is crucial to deliver into the brain target sufficient intensity, which depends on the degree of the ultrasound dephasing by the skull^27,28^. Therefore, the bulk of previous studies regarding correction for skull aberrations of ultrasound have focused on the compensation for the dephasing, and have achieved adequate phase correction^18,29,30,32–39^.

Crucially, however, reversible transcranial applications have a distinct goal—the delivery of deterministic dose that is high enough to ascertain effectiveness, while, critically, also low enough to ensure safety. We have found that for this specific purpose—i.e., the delivery of a deterministic dose, the correction for the *attenuation* is far more important than the correction for the dephasing (**Fig. 6**).

In this regard, existing approaches provide only limited account of acoustic attenuation by the skull. For instance, albeit image parameters of CT scans can approximate acoustic velocity and thus the dephasing, CT (x-ray) and ultrasonic waves rely on distinct physical principles, which has made it difficult to use CT data to account for the acoustic attenuation^53^ (see^54^ for a meta-analysis of the studies; Fig. 6b in particular). Moreover, existing studies currently do not account for the additional key barriers, which involve coupling media, hair, and entrapped bubbles. These additional barriers are substantial, attenuating the transmitted intensity by ~ 0 – 64 % due to coupling^24,25^, by ~ 20% due to hair^22,23^, and by up to 100% due to a local bubble or air pocket^26^. This study provides a practical (**Fig. 2**), accurate (**Fig. 3**), and safe (**Fig. 4**) method and hardware to compensate for these barriers and thus enable effective and safe, reversible applications of ultrasound-based therapies.

RTT is practical in that it takes into consideration all obstacles within the ultrasound path and requires no CT or MRI scans of the skull and the associated simulations. This is possible thanks to i) the relative comparison between ultrasound propagation through the head and through water ii) by firing each transmitting element sequentially while listening for the signal on a specific receiving element (**Fig. 2**). Conceptually, RTT could be considered a kind of ultrasound computed tomography, which has been applied for studying acoustic properties of soft-tissues^55^, bone^56,57^, and skull and brain imaging^58,59^. Unlike tomography, however, RTT specifically aims to correct for object aberrations, performing scans with the object present and absent, and implementing relative computations. In addition, RTT was specifically developed with the objective to correct for the head with respect to transcranial ultrasound therapies. Notably, given the small dimension of each transducer element, RTT implements a virtual line path between each transmit-receive pair of elements, and thus measures all forms of attenuation (reflection, absorption, and scattering) of all obstacles along this defined path.

Two previous groups used an ultrasound-based through-transmit correction approach to improve ultrasound imaging through the skull^32,33^. There are three key differences. First, these approaches used relatively small ultrasound imaging arrays, which enabled them to assume a constant value of phase that the emitting transducers encounter through the nearer skull portion. These algorithms provided only modest correction for our large, therapeutic ultrasound aperture, as the assumption of constant phase and attenuation in the near skull portion is violated. In comparison, RTT amplitude and phase correction algorithms take into account the variable skull distortions both along the transmitting and the receiving arrays. Second, instead of solving for each element’s aberration separately, RTT optimizes the correction of all elements at once to account for the dependence of each element’s correction on that of the others. Third and finally, the goal of RTT is to deliver into the target a deterministic ultrasound intensity. RTT does not aim to compute the attenuation and speedup values by each segment of the skull. To achieve its goal, RTT operates on relative measurements, i.e., measurements performed through the head versus free field (**Suppl. Fig. 6**). These values are used to optimize the relative values of amplitude and phase emission from each element such that the signals from each element arrive into the target with the appropriate amplitude and phase.

RTT is limited to transducer geometries and sections of the skull for which a through-transmit path can be established. We chose to place transducers parallel over the left and right sides of the skull as 1) there is a through-transmit path with minimal incidence angle to the skull 2) the parietal bone has favorable acoustic properties^19^, and 3) this geometry provides convenient coupling. Deviations from this geometry may incur loss of performance or practicality.

RTT also likely depends on specific phased array geometries. We found the method provided accurate correction for the two geometries tested (**Suppl. Fig. 1**). Both geometries had a focal length of 165 mm, which delivers relatively direct through-transmit beams to the receivers while minimizing the angle of incidence to the skull. The more condensed version of the array provided a more accurate correction likely due a higher density of receivers within each beam and due to application through a sm a compromis Dependi the skull^33^. through a smaller, more uniform area of the skull. The individual element sizes of the phased arrays, 6 x 6 mm, were chosen as a compromise between emitted therapeutic power and treatment envelope.

Depending on hardware, the method may be limited to frequencies below 2 MHz, which can penetrate both sides of the skull^33^. We used a frequency common for transcranial ultrasound therapies, 650 KHz^18^. Lower-frequency operation would decrease attenuation and increase the wavelength, making dephasing less severe, and thus would likely improve the compensation further.

The method was also only tested for subcortical targets in a steering envelope spanning 40 mm right to left, 40 mm anterior posterior, and 30 mm superior to inferior in the RAS coordinate system. Future studies should optimize the correction performance through systematic evaluations of the above factors.

The successful stimulation of nerves following the RTT compensation provides a physiologically-relevant validation of the method and thus serves as a proof of concept. Notably, however, the stimulation of peripheral nerves differs significantly from stimulation of neurons in the brain^60^. Therefore, these findings should be interpreted as a physiological readout of the compensated intensity rather than a direct evidence of stimulation of neurons. Nonetheless, these data do highlight the enormous impact that the skull has on modulation of excitable structures. Moreover, nerves and nerve tracks located in deep brain regions appear to be crucially involved in several disorders of brain function^61^. The ability of RTT and the associated hardware to stimulate nerves at depth is therefore critical for effective future treatments. Ultrasonic stimulation of neurons generally requires lower ultrasound intensities than stimulation of nerves^5,52,62^. Moreover, ultrasonic stimulation of nerves in the brain also appears to require lower stimulation intensities compared to nerves in the periphery^62^. Therefore, the intensities found effective for stimulation of peripheral nerves here (**Fig. 7b**) likely represent an upper bound on the intensities needed to stimulate neural structures in the brain.

The drug release data (**Fig. 8**) show that RTT can be used to achieve comparable levels of drug release as in previous *in-vitro* and *in-vivo* studies^12,13,47^. Crucially, however, this holds only when the human skull is compensated for, which was not considered by these previous studies. Like those previous studies, the present study has the caveat that the approach cannot be applied to the human brain until comprehensive safety data are available^47^. The delivery of the deterministic intensity into specific targets through the skull (**Fig. 3**, **Fig. 5**) and the associated deterministic drug release (**Fig. 8b**), nonetheless, present a critical step toward safe applications in humans.

In summary, we developed and tested an approach that accurately and safely compensates for the severe and unpredictable attenuation and distortions of ultrasound by the skull. This method is expected to unlock the full potential of existing and emerging applications of transcranial focused ultrasound, thus realizing their promise in precise and personalized diagnoses and treatments of the brain.

## Online Methods

### Ultrasonic Hardware

RTT was implemented on two hardware platforms. Both systems used two spherical arrays mounted to a frame such that they were positioned opposite to each other and separated by a distance of 180 mm. The array elements of both systems were made of the PMN-PT material, had a surface area of 6 mm x 6 mm, and operated at a fundamental frequency of 650 kHz.

The first system used two spherically focused arrays (radius of 165 mm; 126 elements; 9 x 14 element grid, inter-element spacing of 0.5 mm). Each array had a height of 55 mm and a width of 86 mm, spanning an area of 47.3 cm^2^.

The second system used larger arrays. Each array had 128 elements (8 x 16 grid) with inter-element spacing of 3 mm on a ellipsoid (*R_x_* = 100 mm, *R_y_* = 120 mm, *R_z_* = 165 mm). Each array had a height of 74 mm and a width of 151 mm, spanning an area of 105.6 cm^2^ (**Suppl. Fig. 1b**).

These transducers delivered ultrasound through the parietal and temporal bones of *ex-vivo* skulls. Specifically for each subject, the transducers were orientated in parallel to the left and right sides of the skull. The transducers were driven by a programmable system (Vantage256, Verasonics).

### Targeting

Targeting with ultrasound rests on emitting ultrasound from each element such that the wavefronts arrive into the defined target at the same time. These values can be calculated by 1) considering the acoustic properties of water 2) measurements using a hydrophone. We used the second approach to maximize the targeting accuracy. In these measurements, each element was driven with 10 cycles of a 650 kHz sine wave with an amplitude of 15 V.

### Correction Methods

The goal of the skull correction is to measure the amplitude attenuation and phase speedup of each beam, and compensate for them, ideally to deliver into a brain target the same intensity as if no skull was present.

The signal emitted from each transducer *i* on its path to specific brain target of interest is attenuated by acoustic obstacles (skull, hair, coupling) by a factor of *A_i_*, and sped up by *τ_i_*. Each factor *A_i_* and *τ_i_* is specific to the position of the target due to its unique path through the skull. The aim of the below correction methods is to estimate these values and compensate for them. The compensation scales the amplitude of each beam by a factor of 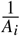 and delays the emission time by *τ_i_*. The measurements are performed in a relativistic way, with the skull present and absent (“free field”, i.e., only water).

#### No Correction

No adjustments to the emission times and amplitudes were performed following the targeting in water.

#### Hydrophone Correction

This correction uses a hydrophone positioned at the target to measure *A_i_* and *τ_i_* directly. These measurements provide the hypothetical ground truth. The relative speed-up time, *τ_i_*, was obtained as the time that maximizes the cross-correlation between the waveform received through the skull and in free field. The relative attenuation, *A_i_*, is measured as the ratio of the peak negative pressure of the two waveforms. The peak negative pressure was computed as the median of the negative cycle peaks over the 10 cycles.

#### Relative through-transmit Correction

In this method, the transducers emitted a 10-cycle, 650 kHz pulse from each of its elements while recording responses from all the other, non-transmitting elements. During the through-transmit scans, the peak pressure amplitude of each transducers was 80 kPa. The entire process of this scan takes less than 1 s to complete. The 650 kHz pulse frequency is the same as that used for the neuromodulation. The equivalence of energy kind (acoustic) and frequency (650 kHz) between the through-transmit measurement and the neuromodulatory ultrasound enables a direct measurement of the ultrasound attenuation and phase shift by the skull and other obstacles in the path, compared to indirect CT imaging methods. This through-transmit measurement is relativistic, performed with the skull present and absent i.e. through water. The relative differences in the received ultrasound waveforms between the two conditions enable the computation of 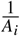 and *τ_i_*. The *A_i_* and *τ_i_* values are computed separately.

The attenuation computation amounts to solving the following system of equations:

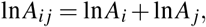

where *A_ij_* is the relative attenuation measured by the through-transmit method for ultrasound propagating from element *j* to element *i*, through both sides of the skull. The attenuation values through the two opposite segments of the skull are multiplicative; hence the logarithmic formulation for attenuation. This linear system of equations can be represented in a matrix form as *Mx* = *b*, where *M* is a matrix of the unitary value coefficients for transmit receive pairs, *x* is a vector of the sought values (*x* = [ln *A*_1_, ln *A*_2_,…, ln *A*_256_]), and *b* is a vector of the measured values (ln *A_ij_*).

This inverse problem can be solved using a variety of methods, including ordinary least squares, truncated singular-value decomposition, and Tikhonov regularization. We applied SVD decomposition on the M matrix, inverted to solve for b, and used Tikhonov regularization to remove high frequency noise as it provided the highest accuracy. The matrix equation M was conditioned in three ways, each of which improves the correction accuracy. First, we selected through-transmit pairs in a target-dependent manner. Specifically, we selected pairs where the angle between the transmitting transducer and the target and the transmitting and receiving transducers was less than or equal to 10°. This angle was chosen as a compromise between maximizing targetable space while minimizing the incidence angle to the skull and thus undesirable beam aberrations. The method was relatively insensitive to this choice; values between 8-15^°^ provided adequate compensation. Formally this selection can be written as *WMx* = *Wb*, where W is a diagonal weighting matrix with diagonal values *w_ij_* for each pair transmit receive elements. Second, we only used information of elements that provided detectable and plausible values (0.01 ≤ *A_ij_* ≤ 0.85). And third, to ensure invertability, we add to the system of equations *Mx* = *b* an additional set of equations that provide initial values for each **x_i_**. These equations are 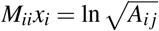, where *M_ii_* = 1 and *j* is the opposing element of *i* with the smallest angle to the target point. The Tikhonov’s regularization parameter, *l*, was set automatically using generalized cross validation^63^. A standard machine running Matlab provides the solution *x* in less than a minute.

The following text describes the algorithm that compensates for the distortion of the ultrasound phase by the skull, so that the ultrasound waveforms from all emitters arrive into a target in phase. The sought phase delays applied to each element are denoted as ***τ*** = [***τ***_1_, ***τ***_2_,…,***τ**_N_*]. Let *s_ij_*(*t*) represent the signal received on a transducer *i* after a brief, 10-cycle pulse is emitted from a transducer *j*. Let the signals received in free field and through the acoustic barriers (skull, hair, coupling, and other barriers) be denoted as 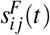 and 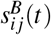, respectively.

The procedure reduces the complexity of all possible combinations of transmit-receive pairs to a simpler case in which a set of transducers focuses onto a single receiver. This way, the speedup experienced by the skull in front of each receiver is a simple linear combination of the through-transmit measurements. Accordingly, the procedure starts by focusing the transmitting elements onto each receiving element in free field. The vector 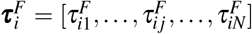 defines the delays in free-field that focus the transmitting signals, 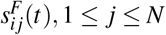, onto an element *i* such that the signals arrive perfectly in phase. Let us define the set of signals received by the element *i* from all transmitting elements as ***s**_i_* = [*s*_*i*1_(*t*), *s*_*i*2_(*t*),…, *s_iN_*(*t*)]. As with the attenuation correction, we control which through-transmit pairs contribute to the equation by including a weighting function ***w**_i_*(*θ, p*) = [*w*_*i*1_, *w*_*i*2_,…,*w_iN_*], where *p* is the target location and *θ* is the acceptance angle of the angle between transducer element *i* and the target and transducer element *i* and the receive transducer. We set *θ* = 10° and let *w_ij_* = 1 for elements with angle below the acceptance angle and *w_ij_* = 0 otherwise, using the same argument as with the attenuation. Let the sum of the received signals, for any vector of delays ***τ***, and any set of weights, ***w**_i_*, be denoted as 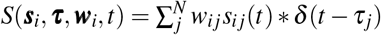.

The goal is to find the delays ***τ*** that account for the speed-up through the obstacles (skull) relative to free-field (water). The focused signal received by element *i* in water, 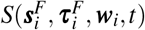, should be delayed by *τ_i_* compared with the signal received through through the skull after applying delays ***τ*** to all other transmitting elements to compensate for their respective speedup due to the skull, i.e., 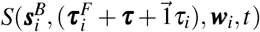. The delays, ***τ***, are found by maximizing the coherence between the through-water and through-skull signals. This amounts to maximizing the following criterion:

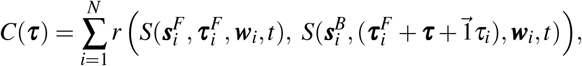

where *r* denotes the Pearson’s correlation.

This optimization problem can be effectively solved through an iterative approach. Starting with 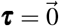 for *n* = 0, we estimate *τ_i_* such as to maximize the correlation 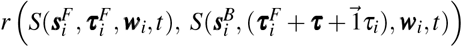. After iterating through each individual element, the delay for each is set to the currently estimated *τ_i_*. The process is then repeated. This iterative procedure terminates when the criterion |*C*| converges to |*C*(***τ***^*n*+1^) - *C*(***τ***^*n*^)| < 0.1 or after 10 iterations. This stopping rule provides a favorable trade-off between compensation accuracy and computation time. For example, a standard machine running Matlab provides a solution in approximately 4 minutes for this stopping rule.

Notably, the keeping track of all 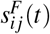 and 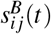 and optimizing the above criterion is necessary to avoid attempts to make exact predictions of phase which are vulnerable to cycle skipping. In cycle skipping, signals that are seemingly in phase are in fact off by a multiple of the period. This could occur if the phase shifts between the free-field and through-skull measurements were computed directly. The method instead attempts to predict time offsets such that waves arrive at the target in phase, regardless if they are off by an integer multiple of the period. **Suppl. Fig. 6** shows raw values of the the attenuation and phase.

We also tested how RTT performs when the correction is performed without a regard to a specific target. To do so, the weight vector is set to include phase information of all elements. The amplitude correction in this case still uses a 10 degree acceptance angle to exclude elements, but now around the normal vector of each element. The correction is calculated once and applied to all targets. The result is shown in **Suppl. Fig. 7**.

### Skulls

Eight *ex-vivo* human skulls were used in this study. The skulls were obtained from Skulls Unlimited (Oklahoma City, OK). The supplier provides ex-vivo specimens specifically for research under a research agreement. A large opening was made at the bottom of each skull to enable field measurements inside the skull. Each skull was degassed overnight in deionized water^20^. Following the degassing at −25 mmHg, the skull was transferred, within the degassed water, into an experimental tank filled with continuously degassed water (AIMS III system with AQUAS-10 Water Conditioner, Onda).

### Hydrophone Field Scans

A capsule hydrophone (HGL-0200, Onda) secured to 3-degree-of-freedom programmable translation system (Aims III, Onda) was used to record the ultrasound field emitted from each element. The hydrophone has a sensitivity of −266 dB relative to 1 V/*μ*Pa and aperture size of 200 *μ*m. This aperture size is well within the ultrasound wavelength (2.3 mm). The 3D field measurements use a step size of 0.2 mm to provide high spatial resolution of each element’s contribution to the total field. The hydrophone scans traversed a volume of 5 x 5 x 5 mm for each target. We also performed wider, 10 x 10 mm planar (XY and YZ) scans for each target. Slices through all fields are shown in **Suppl. Fig. 2**.

The scans were performed both in free-field and through each ex-vivo skull. At each location in the scans, elements were fired individually, and the received signals recorded. Since ultrasound pressure is additive, the total pressure was computed as the sum of the individual constituents. We measured spatial peak intensity of the entire field at each target by taking the maximum intensity value in the measured volume. Position error of this peak and focal volume are quantified in **Suppl. Fig. 5**. For the nerve stimulation (see below), we fired all elements simultaneously as will be the case during applications to the brain.

### RTT in Human Subjects

The hardware and approach described in this article was considered non-significant risk by the Institutional Review Board of University of Utah and approved to be applied in healthy individuals (Protocol #00127033) and patients with major depression (Protocol #00148802). All subjects provided informed consent. Participants were four healthy subjects (4 males, aged between 25-40 years; data points 1-4 in **Fig. 4**) and one subject with major depression (female, 35 years; data point 5 in **Fig. 4**). No hair shaving was necessary as RTT takes hair and other obstacles within the ultrasound path into account. No subject was excluded.

Each subject had the two phased array transducers placed parallel over the left and right sides of their head. The hair length ranged approximately from 1 cm to 4 cm across the subjects. Coupling was mediated using a hydrogel^49^. Standard ultrasound coupling gel was applied to the interfaces between the transducer and the hydrogel, and the hydrogel and the head. The application of the ultrasound gel was not critical given the presence of the hydrogel but improved transmission approximately by a factor of 2. The RTT scan was performed through the entire system of head and coupling. One healthy subject underwent 20 consecutive RTT scans to validate the reproducibility of the measurements (**Fig. 4** shows the first scan). The subject with major depression underwent 3 consecutive RTT scans (**Fig. 4** shows the first scan). No subject reported detrimental effects during or following the procedure.

### Peripheral Nerve Stimulation

Eleven subjects participated in the stimulation (3 females, 8 males, aged between 21-39 years). The study was approved by the Institutional Review Board of the University of Utah (Protocol #00130036). All subjects provided informed consent. No subject was excluded. Subjects placed their thumb into the central location of an *ex-vivo* skull. A 3D-printed positioner held the subjects’ thumb in place. The positioner consisted of a rectangular slot 24 mm wide with a 3 mm semicircular groove that constrained the thumbs lateral, axial, and elevational movement to within 3 mm in each direction. The stimulation was performed inside an ultrasound tank filled with continuously degassed water (AIMS III system with AQUAS-10 Water Conditioner, Onda). Since the ultrasound driving electronics emitted sound and light when stimulating, subjects wore noise-cancelling earmuffs (X4A, 3M; noise reduction rating of 27 dB) and had their eyes closed. Subjects could not hear or see a scheduled stimulus.

Each subject experienced eight distinct stimuli, presented randomly. The ultrasound stimuli were 300 ms in duration. There were three different correction methods—hydrophone correction, no correction, and RTT correction (see above)—and a sham condition. In the sham condition, which was specifically presented for the ideal, hydrophone correction, the ultrasound was programmatically steered 10 mm below the target. For hydrophone- and RTT-corrected stimuli, we varied the intended peak pressure levels across 1.3, 1.55, and 1.8 MPa. The no correction and sham stimuli were tested only at the highest intended pressure level. We performed 10 repetitions of each stimuli producing a total of 80 trials per subject. The stimuli were delivered every 8-12 seconds and randomized so that subjects could not anticipate their onset or type. The subjects were instructed to report a percept with a verbal command (Pain, Vibration, or Tap). A blinded experimenter recorded the percept associated with each stimulus. The response frequency was computed as the proportion of trials in which a nociceptive response was registered.

### Local Drug Release

We manufactured ultrasound-sensitive nanoparticles according to previous approaches^12,13,46^. The procedure specifically followed that of ^47^. As in that study, we used perfluorooctylbromide core given its relatively high stability^47^. As previously, we encapsulated in the nanoparticles the neuromodulatory drug propofol^12,13^. The drug was encapsulated at a concentration of 62.9 *μ*g/mL and a total mass of 12.6 *μ*g. Freshly prepared nanoparticle emulsions were introduced into vials of 8 mm in diameter (1.5 ml polypropylene microcentrifuge tubes, Globe Scientific) at a volume of 0.2 ml, reaching a height of 7 mm. For each ultrasound-release condition (*n* = 10 vials), a vial was placed into a central location of an *ex-vivo* skull, similarly as during the nerve stimulation (see paragraph above). A 3D-printed positioner held the vial in place. The drug release used the same ultrasound conditions, timing, and pressure levels as during the nerve stimulation (see above). The only difference was the exposure time—while the stimulation used 300 ms total time per repetition, the drug release used a total exposure time of 6 s and a single repetition. Specifically, the ultrasound was delivered into the target for 100 ms every 1 s for 60 s. As in the neuromodulation experiment, the no correction and sham stimuli were tested only at the highest pressure level. Propofol released from the nanoparticle emulsions was extracted into a 0.1 mL layer of hexane, as in previous studies^47,64^. The concentration of propofol encapsulated then extracted was quantified using UV-Vis spectrophotometry (Nanodrop 2000, Thermo Scientific). We included an additional condition in which no ultrasound was applied to obtain a baseline for diffusion of propofol into hexane.

## Acknowledgements

This work was supported by the NIH grants R00NS100986, RF1NS128569, and the University of Utah College of Engineering seed grant. We thank Drs. Brian Mickey, Richard Rabbitt, Douglas Christensen and Dennis Parker for helpful comments. We thank John Rolston for the assessment of a skull with a potential hyperostosis. The method and hardware described herein are subjects of a provisional patent.

