## Supplementary Material for "Controlled delivery of ultrasound through the head for effective and safe therapies of the brain"

### Supplementary Material: Controlled delivery of ultrasound into the brain

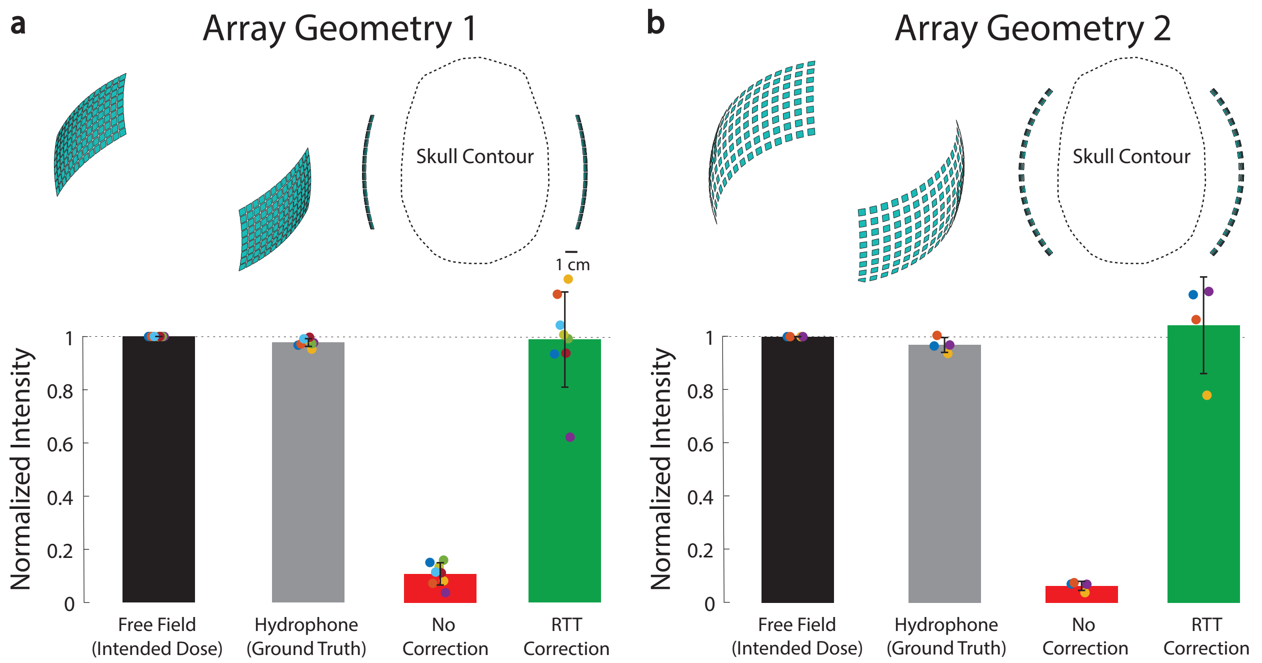


**Supplementary Figure 1. RTT is robust across distinct hardware.**
**a,** Array geometry used in the main text. The associated correction values are the same data as in **Fig.** 3. The correction performed relatively poorly in subject 8 (purple marker), which had the thickest skull with bone outgrowths and possible hyperostosis.
**b,** Array geometry with a larger aperture and the associated correction data.
See Methods for the specific dimensions.

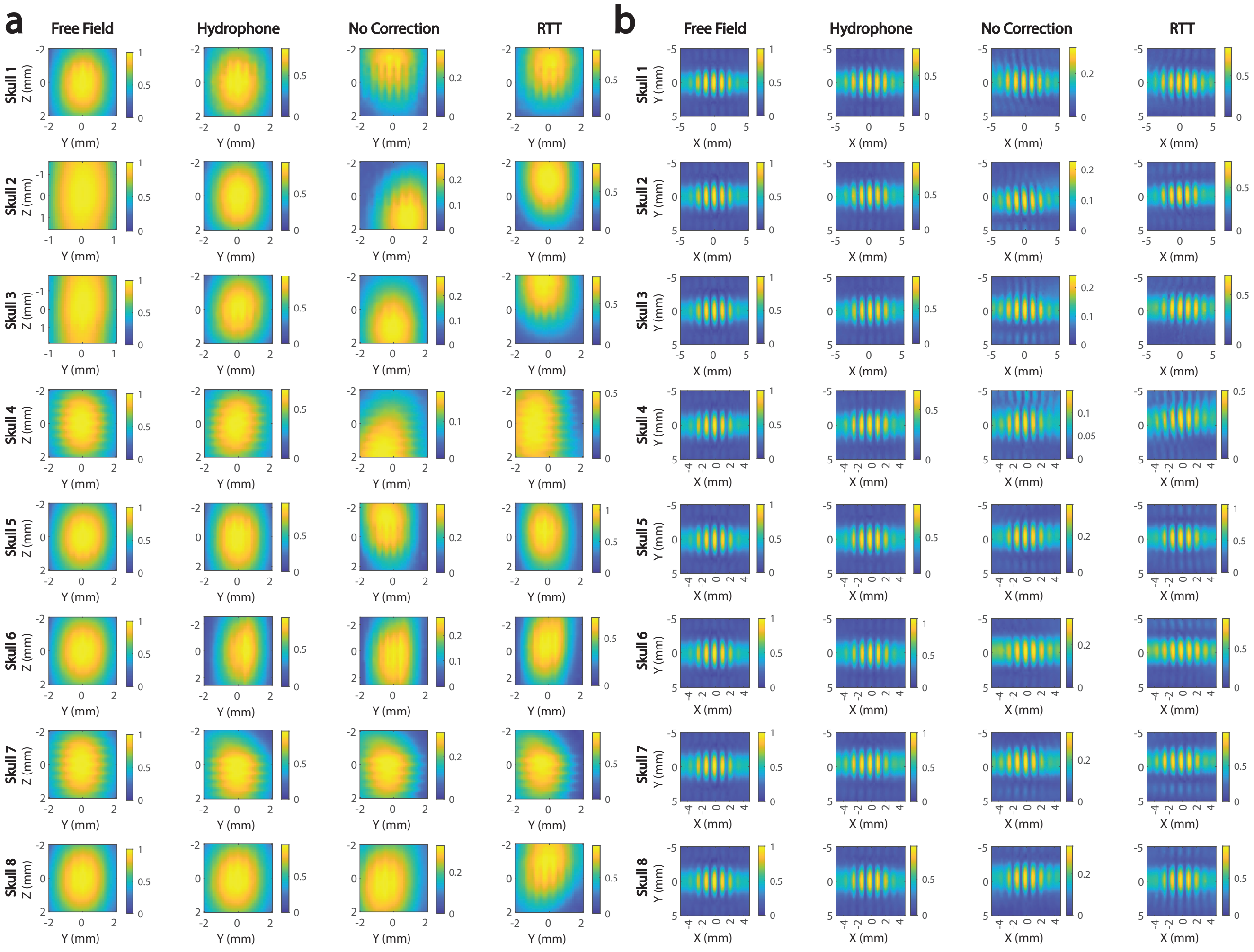


**Supplementary Figure 2.** **Ultrasound pressure fields at central target for all corrections.**

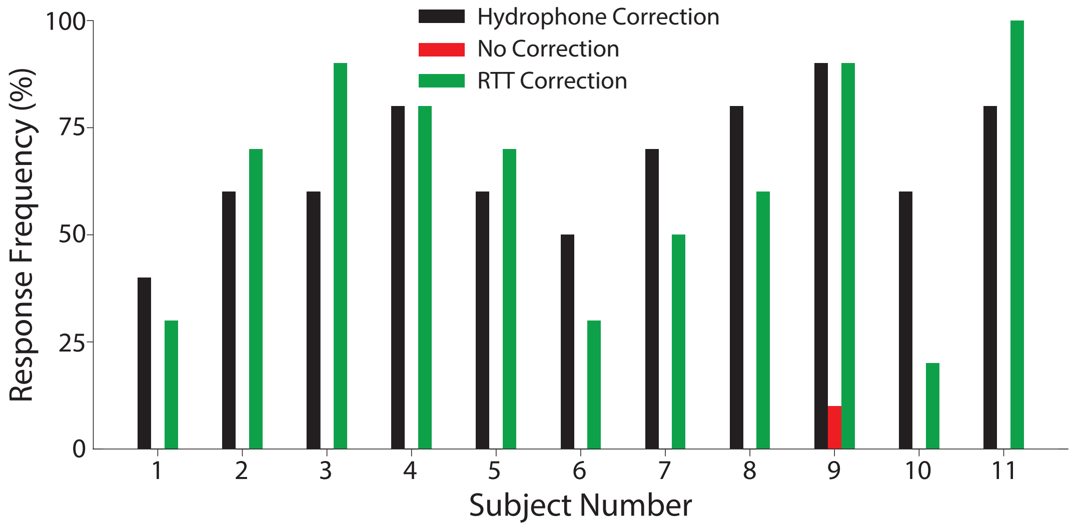


**Supplementary Figure 3.** **Nerve stimulation in each subject.**
Mean response rates for the ideal (black) and RTT (green) correction for each individual subject. When RTT was not applied, there was no significant ultrasonic nerve stimulation (red; $t_{11} = 1.00$, $p = 0.34$, one-sample t-test).

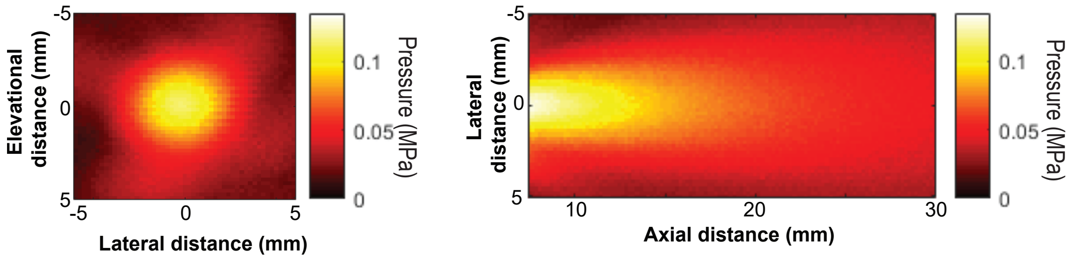


**Supplementary Figure 4. Pressure field produced by each individual element.**
Pressure field of single element of transducer phased arrays measured with hydrophone field scan. Elevational-Lateral plane was taken at an axial distance of 7.5 mm from the transducer face.

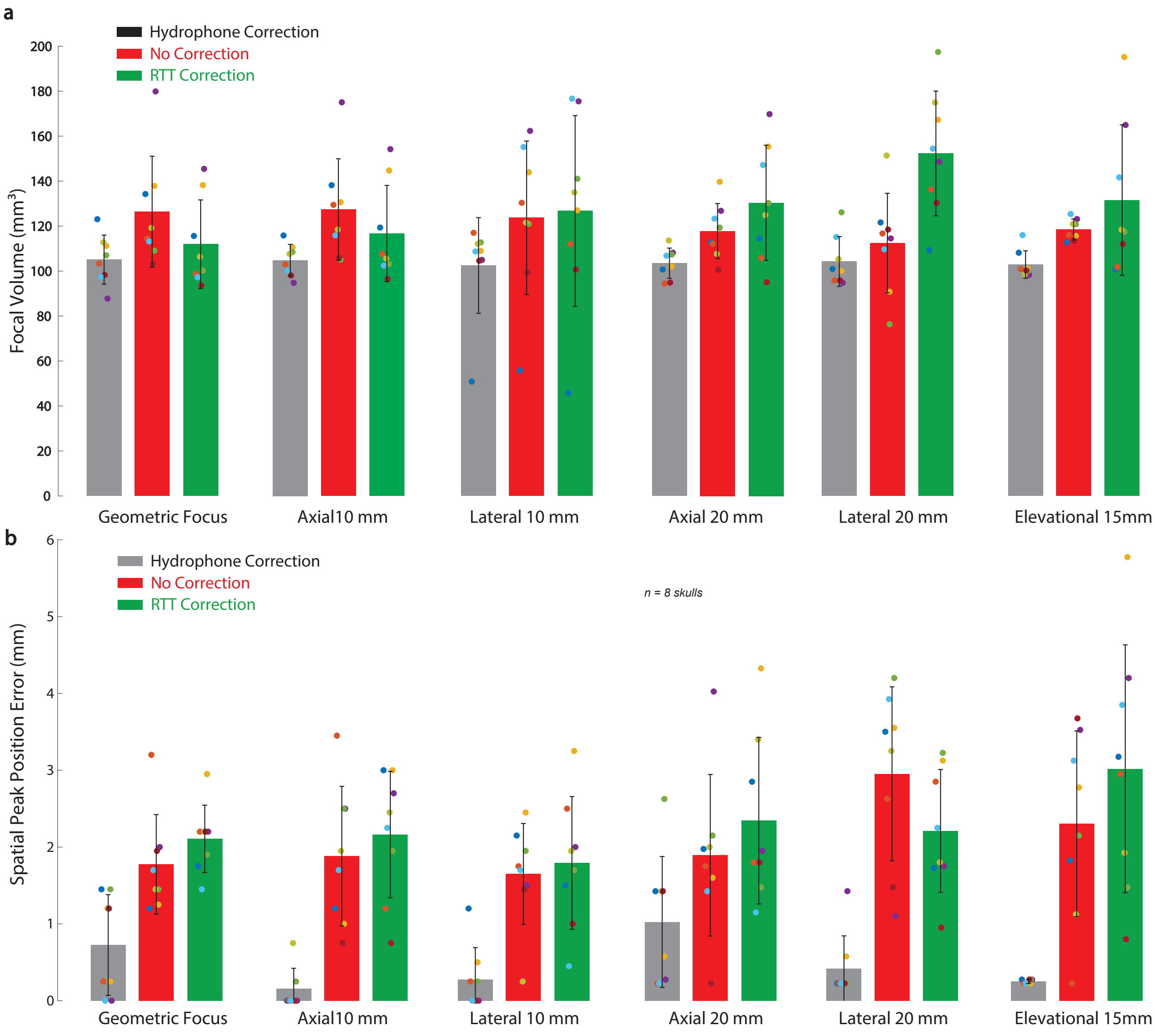


**Supplementary Figure 5. Field volume and target positioning error for each correction.**

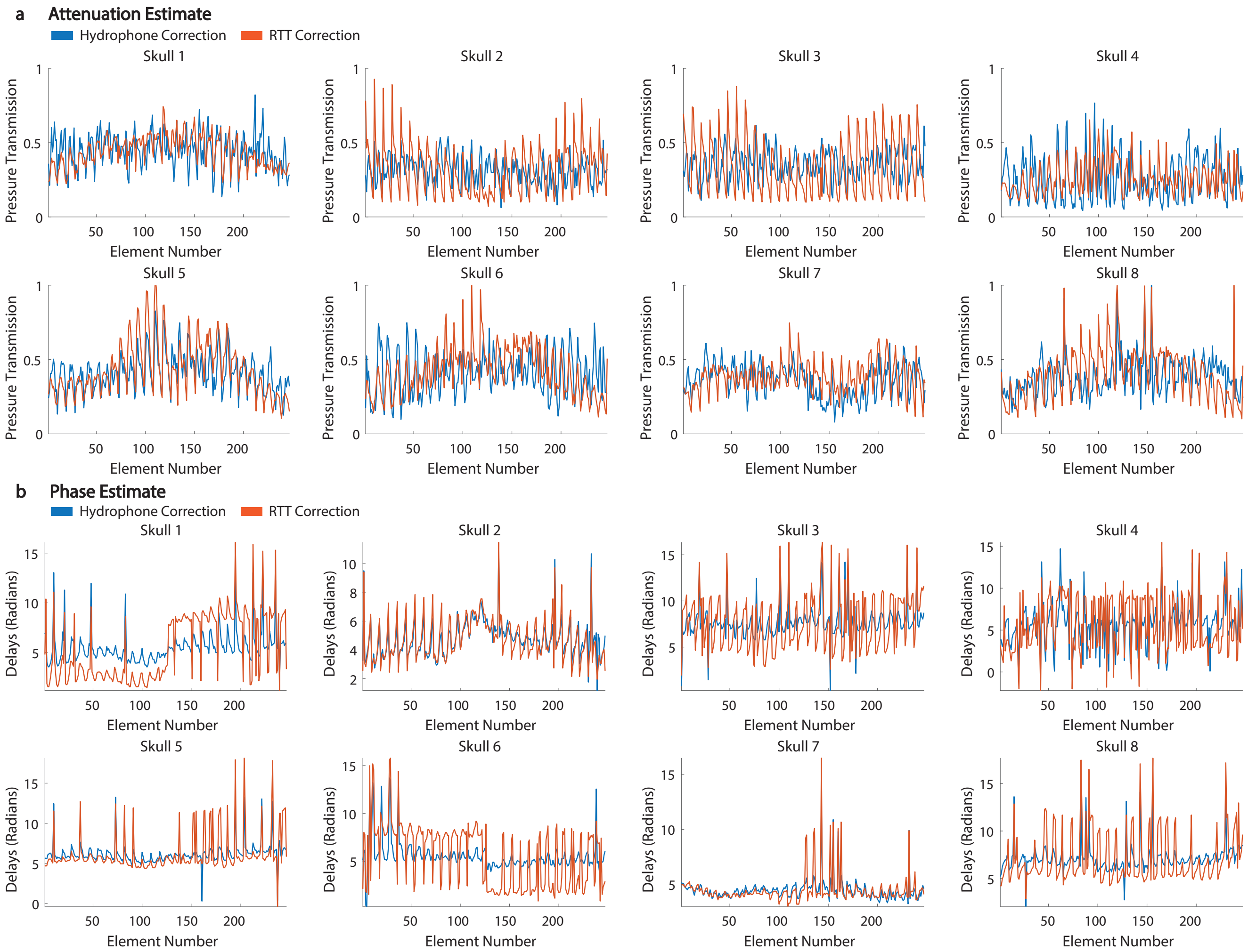


**Supplementary Figure 6. Calculated attenuation and speedups compared to hydrophone.**
The primary goal of the correction is to restore the intended intensity at focus and not to specifically determine the exact attenuation and speedup values through specific sections of the skull. For example, many sets of $A$ and $\tau$ can be combined such that the summed intensity at focus is equal to that of the intended intensity. Nonetheless, the figure shows that the relative values determined by RTT correlate well with those measured by the hydrophone.
**a,** Values of attenuation determined by hydrophone vs values chosen by RTT for each skull.
**b,** Values of phase delays chosen by the hydrophone vs values chosen by RTT for each skull.

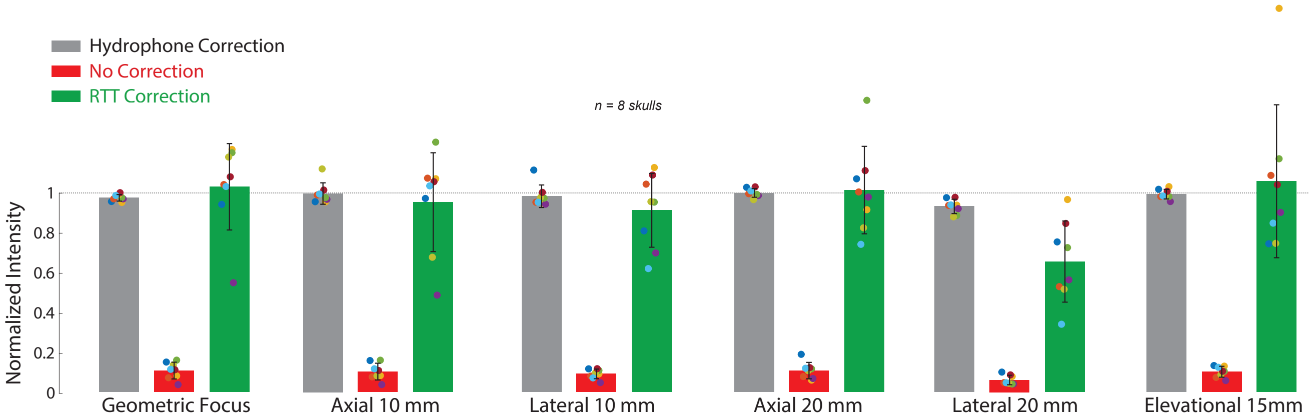


**Supplementary Figure 7. RTT correction using all through-transmit pairs, regardless of the target.** Correction results for no target-dependent weighting (see Methods). In this case, the correction is target-independent. The correction is calculated once, and then applied to any target location. The correction still uses a 10 degree acceptance angle to exclude elements that do not contribute with substantial information.

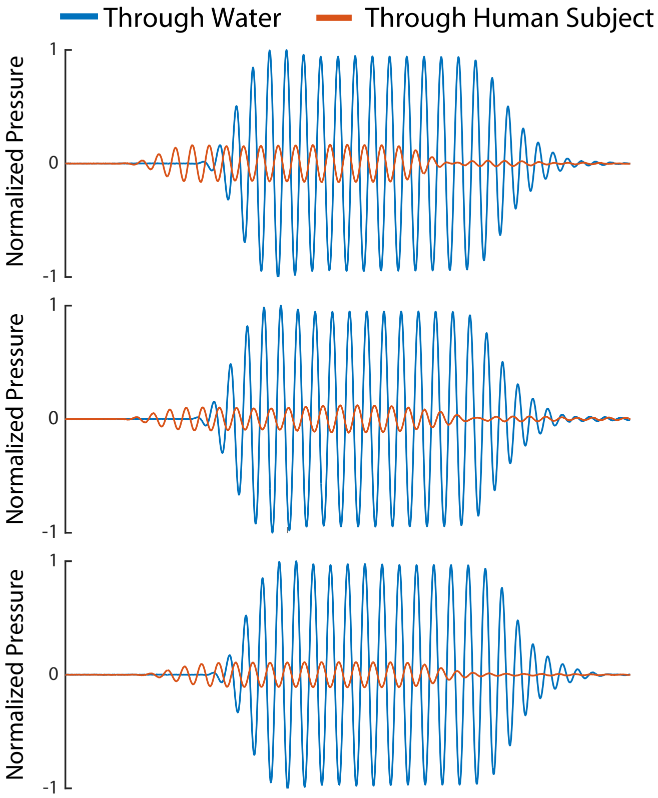


**Supplementary Figure 8. Example through-transmit signals recorded in human subjects with hair.** Received waveforms on three separate transducer elements in water (blue) and after the wave has passed through the head of a human subject (orange).
